# Gains of 12p13.31 delay WNT-mediated initiation of hPSC differentiation and promote residual pluripotency in a cell cycle dependent manner

**DOI:** 10.1101/2021.05.22.445238

**Authors:** Alexander Keller, Yingnan Lei, Nuša Krivec, Edouard Couvreu de Deckersberg, Dominika Dziedzicka, Christina Markouli, Karen Sermon, Mieke Geens, Claudia Spits

## Abstract

Though gains of chromosome 12p13.31 are highly recurrent in hPSC, their impact on differentiation is poorly understood. We identify a reduction in differentiation capacity towards all three germ layers and a subpopulation of residual pluripotent cells that appear during hepatic specification. These cells form as a result of the overexpression of *NANOG* and *GDF3,* whereby NANOG as the primary driver delays activation of WNT signaling, partly as a result of a direct physical interaction with TCF7. Entry into the residual state is determined by cell cycle position at the onset of differentiation and is maintained by a feedback loop between *NANOG* and *GDF3.* These findings highlight the ability of genetically abnormal hPSC to escape correct differentiation and to form residual pluripotent cells, an important risk in the safe clinical translation of hPSC. Our results further refine the molecular mechanisms that underpin the exit from pluripotency and onset of differentiation.

## Introduction

Human pluripotent stem cells (hPSC) acquire recurrent copy number variations (CNVs) as a result of *in vitro* culture (International Stem Cell Initiative *et al.*, 2011; Baker *et al.*, 2016). These CNVs quickly take over the cell culture due to selective advantages such as reduced sensitivity to apoptosis (Avery *et al.*, 2013; Nguyen *et al.*, 2014) or increased proliferation (Ben-David *et al.*, 2014), recently reviewed in Halliwell, Barbaric and Andrews, 2020. However, the impact these CNVs have on the differentiation capacity and tumorigenic potential of the cells is still largely unknown. A number of studies have demonstrated altered differentiation outcomes in karyotypically abnormal lines (reviewed in (Keller *et al.*, 2018)), though few studies to date have systematically evaluated these effects nor identified the mechanisms by which the CNVs influence differentiation. The first such study evaluated the highly recurrent 20q11.21 duplication and found that it impairs neuroectoderm differentiation as a result of altered TGF-β signaling (Markouli *et al.*, 2019), and a recent report has further implicated this gain in altered spontaneous differentiation (Jo *et al.*, 2020).

Another exceedingly common CNV in hPSC is the gain of chromosome 12, which can occur as a trisomy or as a segmental gain at 12p13.31. Previous work has demonstrated that these cells proliferate at a higher rate, in part due to the overexpression of *NANOG* (Ben-David *et al.*, 2014), which also drives significant differences in gene expression in these cells. Gains of chromosome 12 have also been associated with reduced differentiation potential and can result in the presence of undifferentiated cells within teratomas (Werbowetski-Ogilvie *et al.*, 2009; Ben-David *et al.*, 2014). Furthermore, gains of chromosome 12 are associated with germ cell tumors (Atkin and Baker, 1982; Rodriguez *et al.*, 2003; Korkola *et al.*, 2006) and the malignant counterparts of hPSC, embryonal carcinoma cells (Draper *et al.*, 2004; Andrews *et al.*, 2005). However, an in-depth study of the differentiation capacity of hPSC with gains of 12p13.31 is currently lacking.

In this study, we found that segmental gains of 12p13.31 decrease the differentiation efficiency of hPSC and result in foci of residual undifferentiated cells specifically upon hepatic differentiation. We found that this was driven primarily by the overexpression of *NANOG,* which directly interferes with the activation of the WNT-signaling pathway and is modulated by the cell’s position in the cell cycle.

## Results

### Gain of 12p13.31 impairs exit of pluripotency in a subpopulation of hPSC during hepatic differentiation

This study includes three human embryonic stem cell lines containing segmental gains of 12p (hPSC^12p^), VUB14^12p^, VUB17^12p^ and VUB19^12p^, matched to their isogenic genetically balanced counterparts (hPSC^wt^), VUB14^wt^, VUB17^wt^ and VUB19^wt^. The gains of 12p span a 400kb minimal common region of gain, including the pluripotency-associated genes *NANOG* and *GDF3* (Fig. 1A and Fig. S1A). The increased gene copy-number translates into higher mRNA levels of both genes, which were increased by 1.8-fold and 5.6-fold respectively in hPSC^12p^ compared to hPSC^wt^ (p≤0.05 unpaired t test, Fig 1B). We evaluated the tri-lineage differentiation capacity of these lines by inducing differentiation of the cells into hepatoblasts, neuroectoderm and myoblasts, and evaluating the mRNA levels of *HNF4A, PAX6* and *PAX3* expression respectively (Fig. 1C). All three hPSC^12p^ lines displayed an average of 3-fold lower levels of *HNF4A, PAX6* and *PAX3* (p≤0.05 unpaired t-test) relative to their isogenic counterparts, indicating a generally decreased differentiation efficiency. Interestingly, the impaired differentiation was accompanied by high levels of *OCT4* mRNA in hPSC^12p^ hepatoblasts, but not in neuroectoderm or myoblasts (Fig. 1D). To establish if this was an overall low-level residual *OCT4* expression across all cells, or due to a subpopulation of pluripotent cells, we evaluated a full dish of differentiated cells by immunostaining. We identified a substantial subpopulation of OCT4^+^ cells clustered within the HNF4A^+^ differentiated population (Fig. 1E and Fig. S1B), taking a dense colony-like morphology. These OCT4^+^ cells could be isolated, re-expanded in standard hESC culture conditions and re-differentiated, yielding the same mixed HNF4A^+^/OCT4^+^ population after differentiation, suggesting that these cells were pluripotent stem cells (data not shown). To determine if this was a temporary state of pluripotency that might collapse over time, we extended the hepatic progenitor differentiation to 22 days, from which we were again able to isolate pluripotent cells with classic hESC morphology (Fig. S1C). Mouse embryonic stem cells are able to enter into a state of dormancy resembling diapause, which may lay at the basis for residual pluripotent cell formation (Bulut-Karslioglu *et al.*, 2016; Scognamiglio *et al.*, 2016; Ikeda and Toyoshima, 2017). To determine if a similar state of dormancy might occur in hPSC^12p^, we pulse-labeled VUB14^12p^ with EdU at day 8 of hepatoblast differentiation. OCT4^+^ residual pluripotent cells readily incorporated EdU, indicating the cells were actively proliferating and not in a dormant state (Fig. 1F and Fig. S1D).

**Figure 1:**
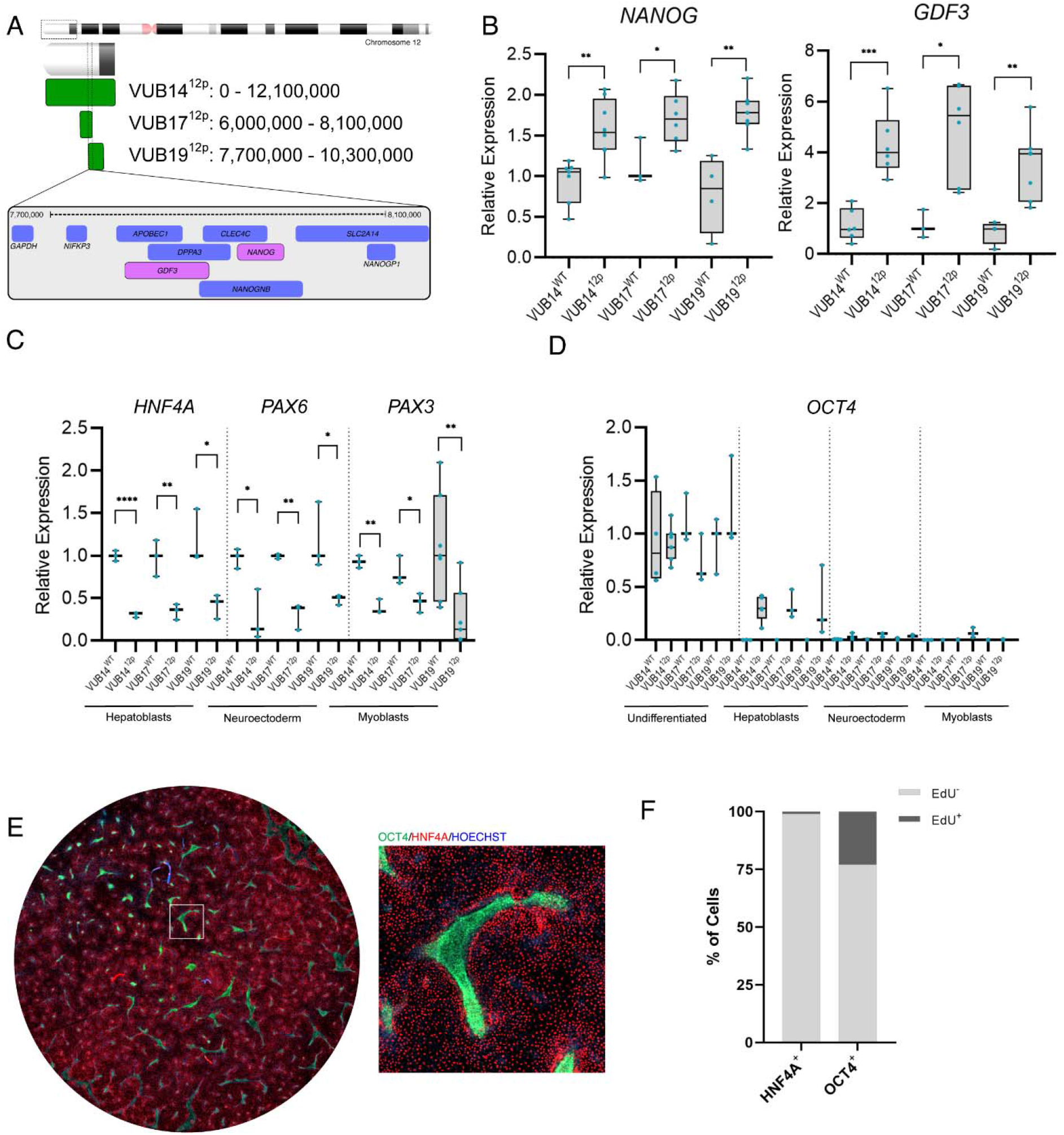
Gain of 12p13.31 impairs exit of pluripotency in a subpopulation of hPSC during hepatoblast differentiation. A) Schematic representation and breakpoints of the minimal overlapping region of gain in VUB14^12p^, VUB17^12p^ and VUB19^12p^. B) Relative mRNA expression of *NANOG* and *GDF3* in hPSC^12p^ and hPSC^wt^. For each isogenic pair, the corresponding undifferentiated hPSC^wt^ was set as reference (n = 3-8). C) Expression of *HNF4A, PAX6* and *PAX3* in hPSC^12p^ and hPSC^wt^ for hepatoblasts, neuroectoderm and myoblasts. For each isogenic pair, the corresponding differentiated hPSC^wt^ was set as reference (n = 3-7). D) Expression of *OCT4* in hPSC^12p^ and hPSC^wt^ for undifferentiated, hepatoblasts, neuroectoderm and myoblasts. For each isogenic pair, the undifferentiated sample was set as reference (n = 3-7). E) Immunofluorescence of 2cm^2^ well of VUB14^12p^ with a magnification of single residual pluripotent colony after hepatoblast induction, OCT4 (green), HNF4A (red), Hoechst (blue). F) Percent of OCT4^+^ residual pluripotent and HNF4A^+^ cells that incorporate EdU after hepatoblast induction, 248/1087 and 15/1721 cells respectively. Statistics: unpaired t-test *= p≤0.05, **= p≤0.01, *** p≤0.001, **** p≤0.0001. See also Figure S1.

### Priming for the residual pluripotent state occurs within the first 12 hours of differentiation

To better understand the differentiation dynamics that lead hPSC^12p^ to form a mixed population upon hepatoblast induction, we performed a time-course single-cell RNAseq. VUB14^12p^ was sequenced in the undifferentiated state and after 12h, 24h, 48h, 72h and 8-days of differentiation. After 8 days of hepatoblast induction, two independent populations can be identified: *OCT4^+^* residual pluripotent cells (Fig. 2A, Cluster 11) and *HNF4A^+^* hepatic progenitors (Fig. 2A, Cluster 12 and 13). The differentiation towards the *HNF4A^+^* hepatic progenitors follows the well-established route through *TBXT*^+^ primitive streak induction at 12h and 24h (Fig. 2A, Clusters 3 and 4) and progression through *SOX17*^+^ definitive endoderm at 48h and 72h (Fig. 2A, Clusters 6 and 8). In the differentiating cells, *OCT4* expression steadily decreases, beginning at definitive endoderm and with a complete loss by day 8 (Fig. 2A and B).

**Figure 2:**
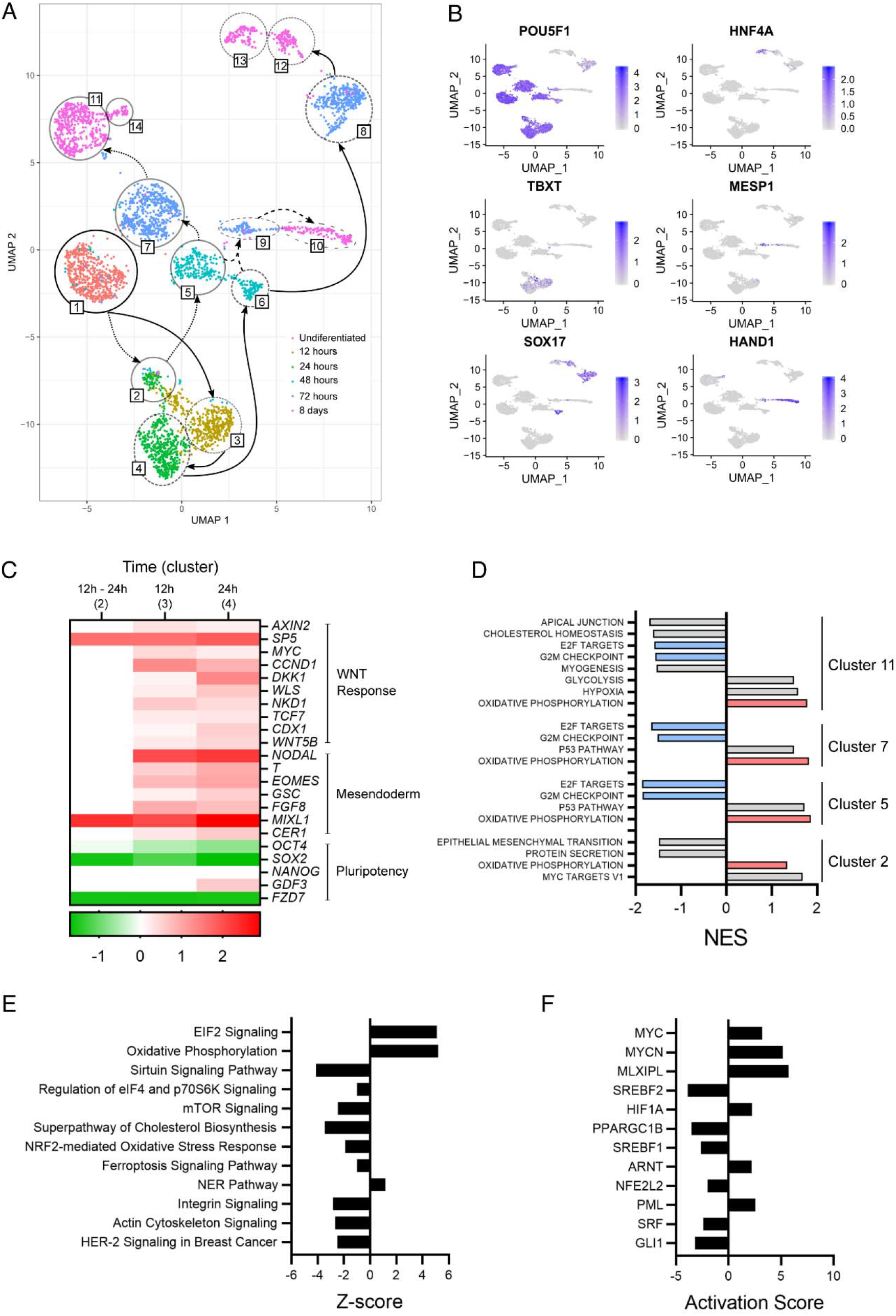
Priming for the residual pluripotent state occurs within the first 12 hours of differentiation. A) UMAP of the single-cell RNA sequencing of undifferentiated VUB14^12p^, and at 12h, 24h, 48h, 72h and day 8 of hepatoblast differentiation. The full line indicates differentiation to hepatoblasts, the dotted lines indicate the route to residual stem cells. The dashed line indicates the route to *HAND1^+^* cells B) Gene expression of pluripotency and differentiation-associated genes in UMAP clusters over the course of 8-day hepatic progenitor induction. C) Heatmap of gene expression of WNT, mesendoderm and pluripotency-associated genes in clusters 2, 3 and 4 relative to cluster 1 (Log_2_FC, FDR: ≤0.0001, white = ns). D) Gene Set Enrichment Analysis using the Hallmark library of clusters 2, 5, 7 and 11 residual pluripotent cells against cluster 1 undifferentiated cells (|NES| > 1, FDR:<0.05). E) Ingenuity Pathway Analysis prediction of differentially activated canonical pathways between cluster 1 and 11, listed in order of significance (|Z-score| > 1, p-value:<0.001). F) Ingenuity Pathway Analysis of predicted differentially activated transcription factors in upstream regulators between cluster 1 and 11, listed in order of significance (|Activation Score| > 2, p-value:<0.001).

At 48h, the lineage of residual pluripotent cells is distinct and marked by the high expression of *OCT4* and no expression of differentiation-associated genes (Fig. 2B, Cluster 5). *OCT4* expression is maintained in these cells through to 72h and 8 days, clearly separating them from their differentiating counterparts (Fig. 2B, Clusters 7 and 11). While residual pluripotent cells form independent clusters at 48h and beyond, *OCT4* levels alone cannot be used to identify them prior to this stage due to its high expression levels in all cells in the first 24h. Interestingly, a cluster comprised of cells from both 12h and 24h forms at the onset of differentiation (Fig. 2A, Cluster 2), independent from the clusters of differentiating cells at 12h and 24h (clusters 3 and 4). 12h and 24h cells in cluster 2 have significantly reduced induction of early mesendoderm and WNT-related genes such as *EOMES, TBXT, NODAL, CCND1* and *DKK1* compared to cells in cluster 3 and 4. However, the WNT response gene *SP5* and mesendoderm marker *MIXL1* are both similarly induced, along with comparable downregulation of the pluripotency-associated genes *SOX2* and *FZD7* (Fig. 2B and C). Cluster 2 cells appear to have begun to respond to the induction of differentiation but have not yet committed to the differentiation program as can be seen in clusters 3 and 4 (Fig 2C). This suggests that cluster 2 cells are the first branching point of the residual pluripotent cells and that formation occurs as early as 12h after the onset of differentiation. These data also indicate that residual pluripotent cells may be forming as a result of poor activation of WNT and mesendoderm genes.

In addition to the *OCT4^+^* and *HNF4A^+^* cells seen at day 8, a third population of *HAND1^+^* cells can be identified (Fig. 2A, Cluster 10), which account for the small population of cells that were neither *OCT4^+^* nor *HNF4A^+^* in the immunofluorescent stains (Fig. 1E and Fig. S1B). While the ancestor cells of the hepatic and residual pluripotent clusters can be traced back to the first 12-24h of differentiation, the *MESP1^+^* ancestor cells (Fig. 2A Cluster 9) that progress into the *HAND1^+^* cluster at day 8 do not appear until 72h after the onset of differentiation (Fig. 2A and B). This suggests that the initial pool of cells that leads to their formation is either cluster 5 or 6 at 48h and not the undifferentiated day 0 cells (cluster 1), or clusters 2, 3 or 4 of 12h and 24h cells. *MESP1^+^* is associated with the posterior primitive streak, which is a precursor to mesoderm fates (Loh *et al.*, 2014), and so would be unlikely to develop directly from *SOX17*^+^ cells that have already committed to endoderm. One possibility is that the residual pluripotent cells act as a pool of cells that continuously differentiate and feed into mis-specified cell fates. In support of this, residual pluripotent cells at day 8 (cluster 11) has a small branch of cells that are also *MESP1^+^ and HAND1^+^* (Cluster 14), indicating a portion of the cells may be in the early stages of misspecification.

### Residual pluripotent cells do not show altered gene expression of pluripotency-associated pathways

Given that the residual pluripotent cells clustered separately we compared cluster 1 undifferentiated cells to cluster 11 residual pluripotent cells in order to determine how the hPSC^12p^ and residual pluripotent state differed. We identified 668 significantly differentially expressed genes between the two groups (|Log_2_FC| >0.5, FDR:<0.001). Gene Set Enrichment Analysis (GSEA) and Ingenuity Pathway Analysis (IPA) analysis show that cluster 11 residual pluripotent cells do not have differential activation of pathways commonly associated with pluripotency such as FGF/ERK, TGFβ or OCT4/SOX2/NANOG suggesting that they are in a similar state of pluripotency to their undifferentiated counterparts. However, both GSEA and IPA predict that residual pluripotent cells have active oxidative phosphorylation (OXPHOS) (Fig. 2D and E). GSEA indicates that OXPHOS was also significantly enriched in residual pluripotent clusters 2, 5 and 7, indicating that the metabolic switch is likely activated early on in the residual pluripotent transition. Also common amongst the residual pluripotent cells after 48h was a negative enrichment for G2M checkpoint and E2F target genes (Fig. 2D). We then looked at the number of significantly differentially expressed genes that drive this gene expression signature. We found that 58 genes drove the prediction of activated OXPHOS, however, only 16 and 20 differentially expressed genes drove the prediction of negative enrichment of G2M checkpoint and E2F targets, respectively (supplemental table 1-3). Interestingly, amongst the top transcriptional regulators predicted to be active by IPA are the cancer-associated genes *MYC, MYCN* (Gabay, Li and Felsher, 2014) and *HIF1A* (Zhong *et al.*, 1999) (Fig. 2F). *HIF1A* is associated with a response to hypoxia, which is suggested to be active by GSEA in cluster 11 residual pluripotent cells (Fig. 2D). These data suggest that residual pluripotent cells are transcriptionally highly similar to the normal pluripotent state and that the main difference between them is the activation of OXPHOS and as well as the cancer associated genes *MYC*, *MYCN* and *HIF1A.*

### Overexpression of NANOG and GDF3 induce entry into residual pluripotent state in hPSC^12p^

The common region of 12p gain spans *NANOG* and *GDF3,* and constitutive overexpression of both genes has been shown to maintain hPSC in an undifferentiated state under differentiation-promoting conditions (Levine and Brivanlou, 2005; Darr, Mayshar and Benvenisty, 2006; Chambers *et al.*, 2007). In the case of *GDF3,* this is likely as a result of its role as a BMP inhibitor and as a NODAL analog when expressed at high levels (Chen *et al.*, 2006; Ariel J. Levine, Levine and Brivanlou, 2009). We therefore hypothesized that the excess of both *NANOG* and *GDF3* plays a role in delaying the exit from pluripotency and inhibiting the initiation of mesendoderm differentiation.

Given that the segregation of the differentiating and non-differentiating cells occurred very early into the differentiation process, we investigated the impact of the over-expression of these genes at the onset of differentiation. We treated VUB14^12p^ with siRNA for either *NANOG* or *GDF3* for 24h prior to the initiation of differentiation and evaluated the cells at the end point, after 8 days of hepatoblast induction. We knocked down the expression to levels that closely matched those seen in the genetically normal isogenic hPSC lines, restoring the expression to close to normal levels (Fig. S2A).

Knockdown of *NANOG* led to the full rescue of differentiation, restoring levels of *HNF4A* to those seen in VUB14^wt^ and completely eliminating the residual undifferentiated cells (Fig. 3A and 3B). In contrast, knockdown of *GDF3* led to the loss of residual pluripotent cells but did not fully rescue the differentiation, instead creating a mixed population of *HNF4A^+^* and mis-specified cells. These results establish that over-expression of both *NANOG* and *GDF3* are necessary to prevent cells from initiating mesendoderm differentiation and subsequently maintain residual pluripotency. *NANOG’s* role in interfering with the onset of mesendoderm has previously been recognized through constitutive over-expression studies (Vallier *et al.*, 2009) and these results suggest a key supporting role for *GDF3.*

**Figure 3.**
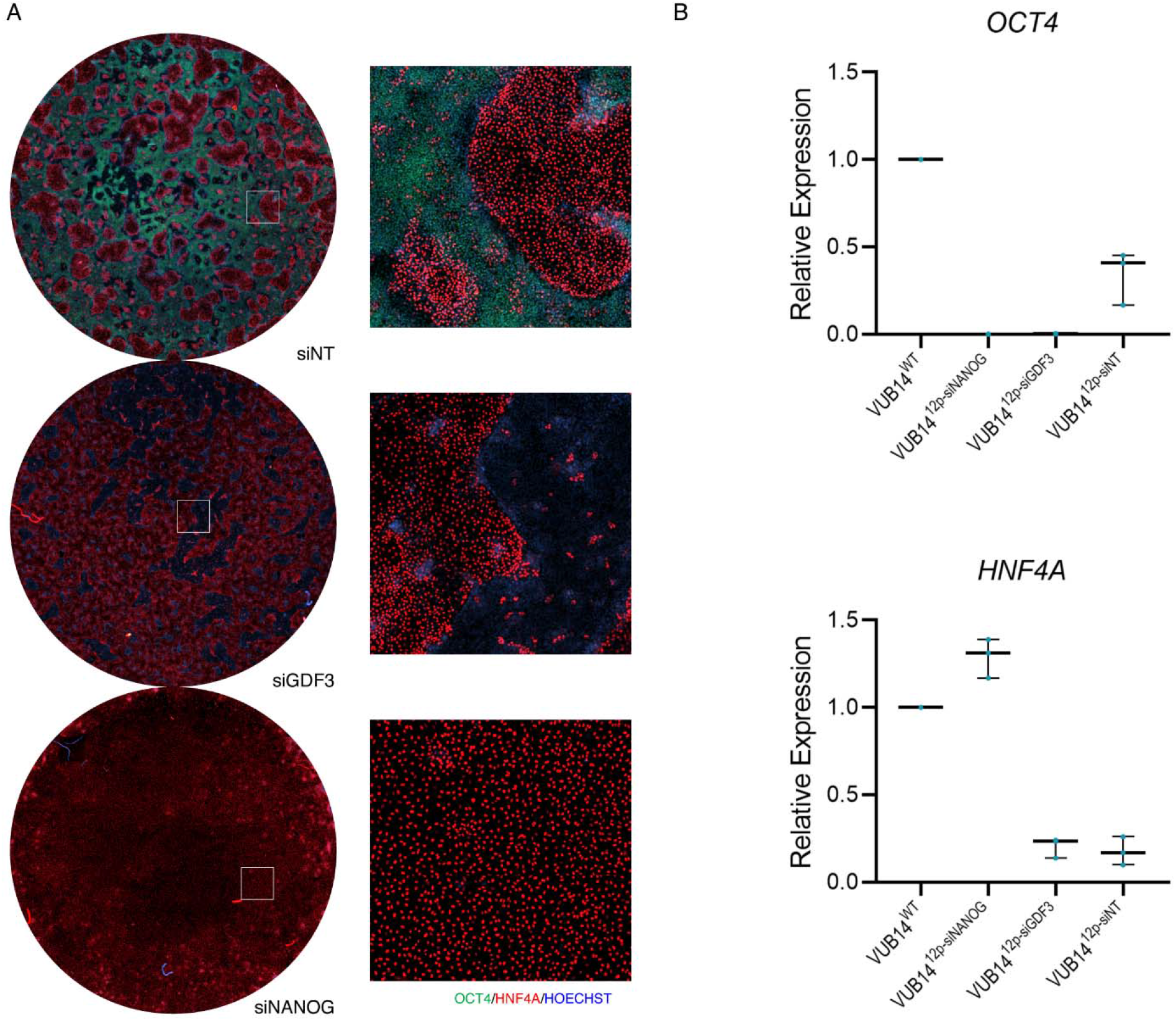
Overexpression of *NANOG* and *GDF3* induce entry into residual pluripotent state in hPSC^12p^. A) OCT4 (green) and HNF4A (red) immunostaining of day 8 hepatoblast of VUB14^12p^ after 24h knock-down of GDF3 and NANOG prior to the onset of differentiation. B) *OCT4* and *HNF4A* gene expression of day 8 hepatoblast of VUB14^12p^ after 24h knock-down of *GDF3* and *NANOG* prior to the onset of differentiation (n = 3). See also Figure S2.

Under normal conditions, endogenous activation of the BMP signaling is necessary for anterior primitive streak specification, the precursor to definitive endoderm and the hepatic lineage. In line with this, differentiating cells at 24h of hepatoblast induction are positive for pSMAD1 (Fig. S2B). We propose that the increased *GDF3* expression in hPSC^12p^ results in the inhibition of the endogenous induction of the BMP signaling during the onset of differentiation, enhancing the effect of *NANOG* in impairing the exit of pluripotency and in subsequently maintaining the cells in the pluripotent state. In support for this mechanism, we found that in hPSC^12p^, the addition of low concentration of BMP4 to bypass the BMP-inhibiting effect of *GDF3* during the first 24h of hepatic progenitor induction, leads to the same outcome as knocking down *GDF3* (Fig. S2C).

Conversely, as the *NANOG* knock-down experiments show, overexpression of *GDF3* alone is not sufficient to impair correct differentiation. In line with previous reports indicating that *GDF3* is a direct target of *NANOG* (Fischer *et al.*, 2010), we found that upon *NANOG* knock-down, the expression of *GDF3* was also reduced, albeit not restored to normal levels (Fig. S2A). This indicates that the 5.7-fold increase in *GDF3* expression in hPSC^12p^ is a result of the compounding effect of its increased copy number and a change in regulation brought upon by *NANOG* overexpression. In the *NANOG* knockdown experiments, the concomitant decrease in *GDF3* expression results in its level to be insufficient to effectively inhibit BMP and to interfere with differentiation.

### hPSC^12p^ have a delayed onset of WNT activation as a result of NANOG overexpression

Next, we aimed at elucidating the mechanisms by which *NANOG* overexpression is driving the residual pluripotent cell formation specifically during hepatoblast induction and not in neuroectoderm nor myoblast induction. We hypothesized that the specific factors involved in the hepatic induction might be key to the residual pluripotent cell formation. To generate hepatoblasts, cells are first specified to definitive endoderm in activin A and the GSK3β inhibitor CHIR99021 containing medium for 24h, followed by 24h in activin A, after which cells are further driven to hepatic progenitors in KO-SR-based hepatoblast medium (adapted from (Cameron *et al.*, 2015)). We cultured VUB14^12p^ in the presence of either activin A or CHIR99021 alone, at the same concentration and timing used for hepatoblast specification and evaluated the differentiation at day 8 for the presence of residual pluripotent cells. Activin A alone yielded a population of mis-specified *HNF4A^-^/OCT4^-^* cells, whereas a significant population of *OCT4^+^* residual pluripotent cells were present when CHIR99021 was administered alone, with only limited expression of *HNF4A* (Fig. 4A and 4B). This indicates that WNT signaling plays an important role in inducing the residual pluripotent cell formation in hPSC^12p^. WNT signaling is known to play a role in both hPSC pluripotency (Sato *et al.*, 2004; Kim *et al.*, 2013), and mesendoderm differentiation (Gadue *et al.*, 2006) based on its concentration and context, with low concentrations and short exposure leading to the temporary maintenance of pluripotency, and higher concentrations and prolonged exposure inducing differentiation (Tsutsui *et al.*, 2011; Singh *et al.*, 2012; Huang *et al.*, 2015). The low induction of WNT-related genes seen in the subpopulation of hPSC^12p^ at 12h and 24h of differentiation (Figure 2A, cluster 2) suggests that hPSC^12p^ have a delayed or decreased response to CHIR99021. As a result, in hPSC^12p^, the 24h WNT signal-induction may induce pluripotency maintenance instead of promoting differentiation.

**Figure 4.**
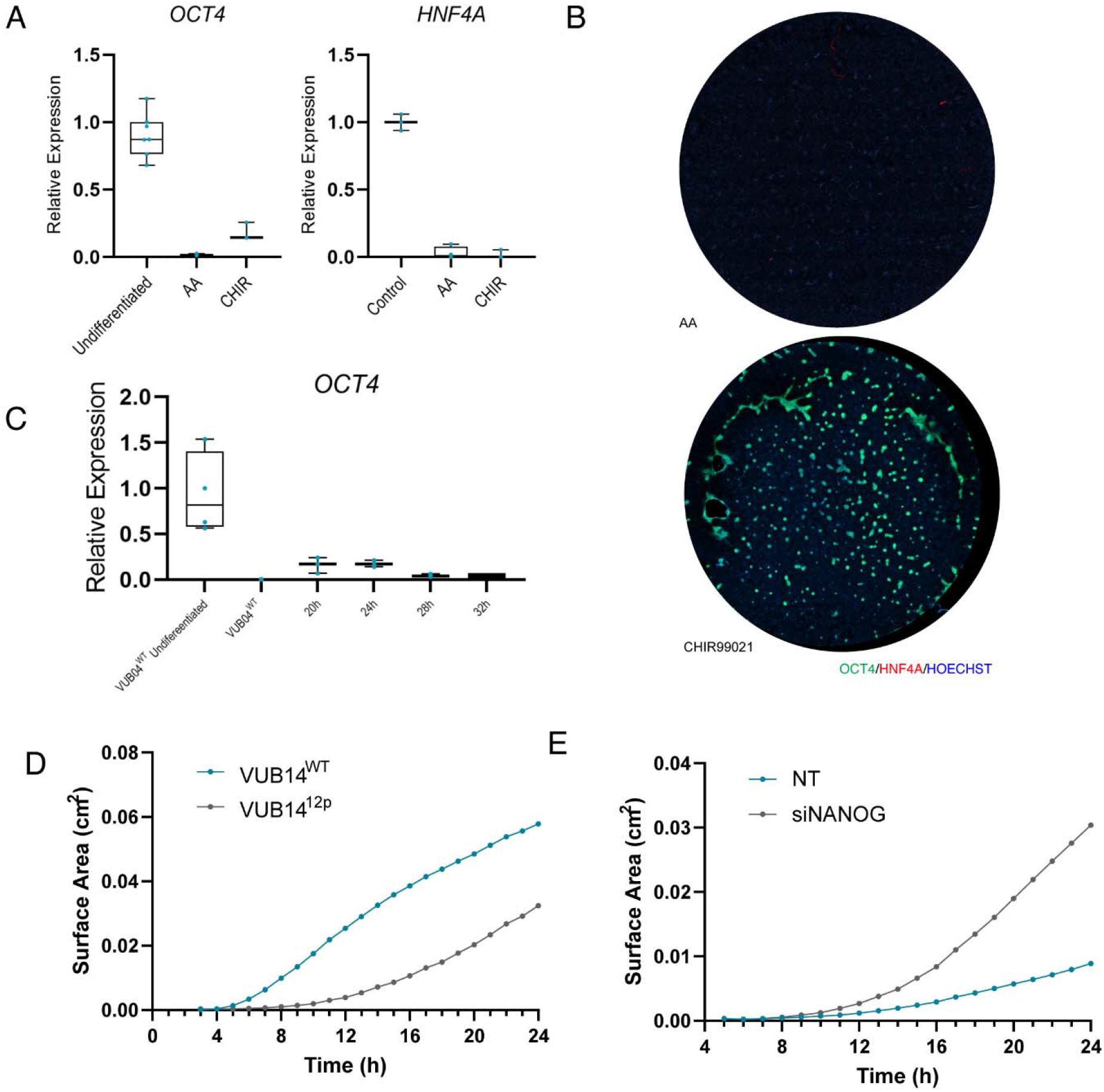
hPSC^12p^ have a delayed onset WNT activation as a result of *NANOG* overexpression: A) Gene expression of *OCT4* and *HNF4A* in day 8 hepatoblasts VUB14^12p^ after induction with only activin A or CHIR99021 (n = 3-4) B) OCT4 (green) and HNF4A (green) immunostaining of day 8 hepatoblasts of VUB14^12p^ after induction with only activin A or CHIR99021 C) *OCT4* gene expression in day 8 hepatoblasts of VUB14^12p^ after extending CHIR99021 treatment in 4h increments (n = 3). D-E) GFP+ surface area in VUB14^WT-TGFP^ and VUB14^12p-TGFP^ during first 24h of hepatic induction under control and *NANOG* knockdown conditions. Values normalized to surface area covered by cells. See also Figure S3

To test this, we exposed VUB14^12p^ and VUB19^12p^ to activin A and CHIR99021 for 20h, 24h, 28h and 32h, followed by 24h of activin A and hepatoblast medium until day 8. As we increased exposure time to 32h, in 4h increments, the proportion of OCT4^+^ residual pluripotent cells seen at day 8 decreased (Fig. 4C & Fig S3A). Interestingly, continued exposure for 48h abrogates all residual pluripotent cells, but also led to near total loss of HNF4A^+^ hepatoblasts by day 8, indicating that the cells are driven towards a different lineage as a result of excessive WNT induction (Fig S3A). Extended WNT signaling is known to induce mesoderm differentiation and it has been reported that during definitive endoderm specification, WNT induction is crucial for 24h, but should be removed or inhibited after this phase to increase specification to anterior primitive streak (Loh *et al.*, 2014). The timing window to specify cells through anterior primitive streak and towards definitive endoderm is therefore shifted in hPSC^12p^ but appropriate factor sequence remains crucial for correct specification.

To further investigate the delayed onset of WNT signaling, we generated a TOP-GFP WNT reporter line of VUB14^WT^ and VUB14^12p^ (VUB14^WT-TGFP^, VUB14^12p-TGFP^) (Fuerer and Nusse, 2010). Live-imaging over the course of 24h showed a greater proportion of VUB14^WT-TGFP^ activating WNT signaling than VUB14^12p-TGFP^ (Fig. 4D and Fig. S3B). Furthermore, siRNA knockdown of *NANOG* in VUB14^12p-TGFP^ led to a greater proportion of GFP^+^ cells after 24h compared to the non-targeting control (Fig. 4E and Fig S3C). It was recently reported that NANOG physically interacts with TCF7, preventing β-catenin from binding to TCF7 in zebrafish (He *et al.*, 2020). Given that the β-catenin – TCF7 interaction is critical for mesendoderm induction (Sun *et al.*, 2017), it is possible that a similar mechanism in hPSC^12p^ could play a role in delaying the onset of differentiation. To test if NANOG and TCF7 interact directly in the human, we transiently overexpressed NANOG and TCF7 in 293T cells and performed coimmunoprecipitation (Co-IP) to determine if they physically interact. We found that NANOG directly binds to TCF7 in the human, likely causing direct interferences between TCF7 and β-catenin (Fig. S3D). Taken together, these results indicate that *NANOG* overexpression attenuates the activation of WNT signaling, in part through direct interference with TCF7, thereby delaying the onset of differentiation.

### The delayed activation of WNT signaling in hPSC^12p^ is cell cycle dependent

The next question we addressed is why the onset of differentiation is only inhibited in a portion of hPSC^12p^. One possibility was that the delayed WNT activation was a stochastic event that affected only a small number of cells that continued to rapidly proliferate up to day 8 of differentiation. To test this we generated an RGB reporter line (Weber *et al.*, 2011; Wu *et al.*, 2016) of VUB19^12p^ (VUB19^12p-RGB^) in which each cell has a random number of inserts of mCherry(Red), Venus (Green) and mTagBFP (Blue). This gives each cell a unique color based on its combination and number of insertions of the three genes, allowing for lineage tracing. We live-imaged VUB19^12p-RGB^ over the course of hepatoblast differentiation (Fig. S4A) and stained for OCT4 at the end of differentiation (Fig. S4B). We found that the colonies of OCT4^+^ cells were comprised of multiple colors, indicating each colony was not of clonal origin (Fig. 5A) and suggesting that a significant part of the starting cells go on to form residual pluripotent cells.

**Figure 5.**
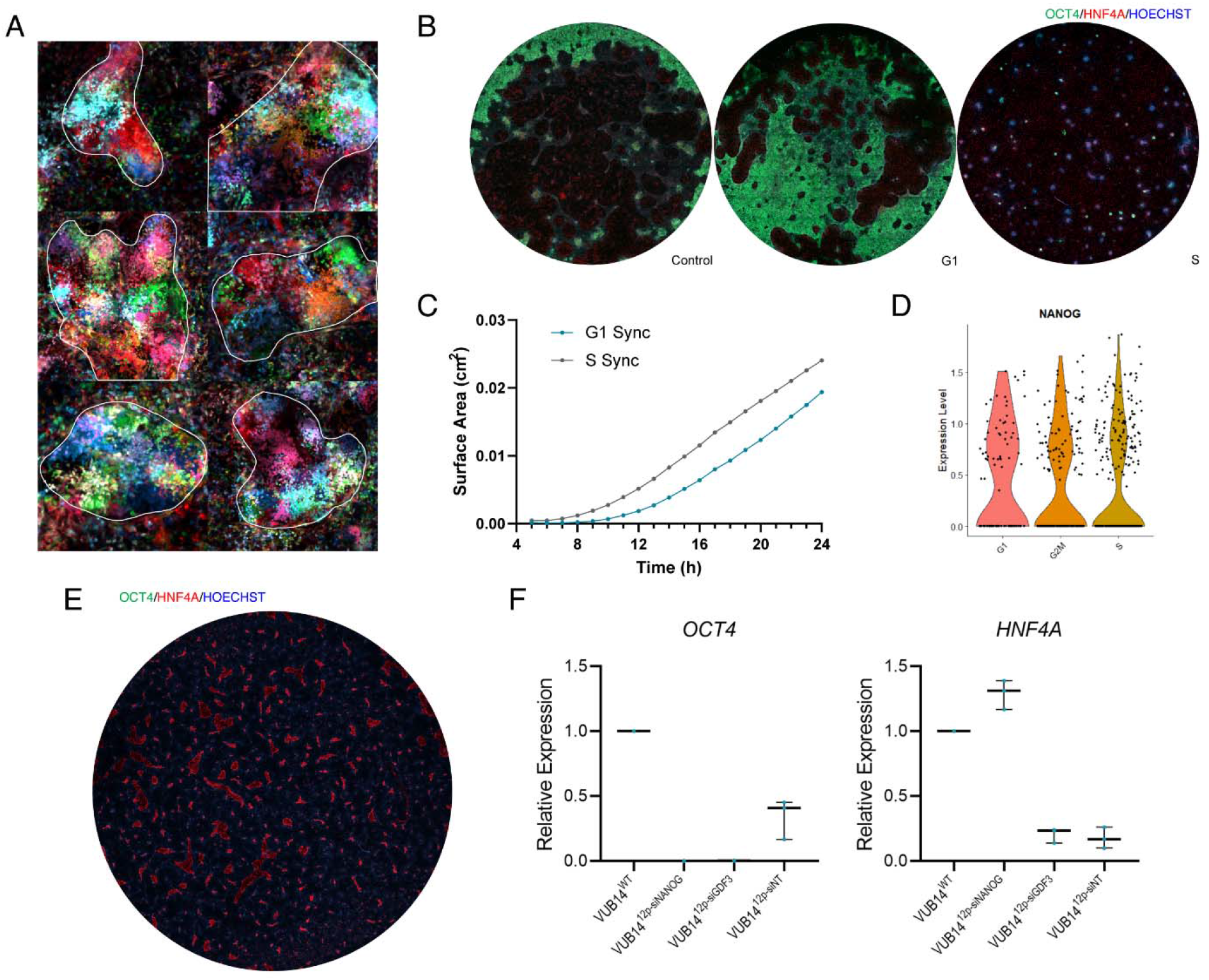
Delayed WNT signaling in hPSC^12p^ is cell cycle dependent: A) Live cell images of residual pluripotent colonies in VUB19^12p-RGB^. B) OCT4 (green) and HNF4A (red) immunofluorescent stains of day 8 hepatoblast of VUB14^12p^ after synchronization to G1 or S. C) GFP+ surface area in VUB14^12p-TGFP^ during first 24h of hepatic induction after synchronization to G1 and S. Values normalized to surface area covered by cells. D) Distribution of *NANOG* expression in single cells in undifferentiated VUB14^12p^ in G1, S and G2M. E) OCT4 (green) and HNF4A (red) immunofluorescent stains of day 8 hepatoblast of VUB14^12p^ after treatment with SB-431542 starting from day 3. F) Gene expression of *OCT4* and *HNF4A* after continuous knockdown of *GDF3* on day 3, 5 and 7 of hepatoblast differentiation (n = 3). See also Figure S4.

Next, we tested if this heterogeneity was the result of each individual cell’s position within the cell cycle at the onset of differentiation. It has been reported that hPSC are most competent to initiate differentiation while in G1 and maintain pluripotency while in S and G2/M (Sela *et al.*, 2012; Pauklin and Vallier, 2013; Gonzales *et al.*, 2015). To test this, we treated VUB14^12p^ with nocodazole for 16h to synchronize the cells to G2/M, and following a 2h and 8h recovery period, induced the differentiation of G1 and S-enriched cells respectively (Yiangou *et al.*, 2019)(Fig. S4C). Surprisingly, we found that hPSC^12p^ enriched to S, but not G1, led to a substantial reduction of residual pluripotent cells and rescue of differentiation (Fig. 5B). In contrast, G1-enriched cells had an increased proportion of residual pluripotent cells relative to the unsynchronized control. To confirm these findings, we synchronized VUB14^12p-TGFP^ to G1 and S and live imaged the cells over the course of 24h. Similarly, we observed WNT activation earlier in S than in G1-enriched cells (Fig. 5C and Fig. S4D). Importantly, a 6h pulse of CHIR99021 in G1 synchronized cells was insufficient to lead to the formation of a significant population of residual pluripotent cells, indicating that there is a minimum activation of the WNT pathway necessary for their formation (data not shown). Lastly, it has previously been demonstrated that cells with a high culture density enrich in G1 (Wu, Fan and Tzanakakis, 2015). Upon initiating differentiation in hPSC^12p^ at a high initial seeding density (>80%), we also observed increased residual pluripotent cells at day 8 of hepatoblast induction and reduced WNT activation at 24h (Fig. S4E).

*NANOG* expression has previously been reported to be heterogenous in hPSC, with distinct populations of *NANOG^hιgh^* and *NANOG^low^* cells (Fischer *et al.*, 2010). To explore the possibility that *NANOG* expression could vary between the different phases of the cell cycle in hPSC^12p^, we split the cells in cluster 1 (Fig. 2A) based on their cell cycle stage (Tirosh *et al.*, 2016). We found that *NANOG* mRNA levels were comparable between the different cell cycle phases (Fig. 5D). Furthermore, we found that the expression of *NANOG* was similarly variable to *SOX2* and *OCT4* in the undifferentiated state, with no clear separation into *NANOG^high^* or *NANOG^low^* populations (Fig. S4F).

These findings strongly suggest that the heterogenous nature of the differentiation is cell-cycle dependent. hPSC^12p^ that are in G1 upon exposure to CHIR99021 have a higher chance to fail to reach the WNT signal threshold required to correctly initiate differentiation, instead remaining behind as residual pluripotent cells. Cells synchronized to S more readily reach this threshold and properly initiate differentiation.

### GDF3 is vital to NANOG’s ability to maintain the undifferentiated state in differentiation-promoting conditions

Finally, we aimed to understand how the residual pluripotent cells were able to maintain the undifferentiated state for extended periods of time, despite the absence of FGF2 or TGFβ, both of which are key to maintain regular hPSC cultures. It has been previously reported that in addition to its role as a BMP inhibitor, GDF3 also acts as a NODAL analog when present at high levels (Ariel J Levine, Levine and Brivanlou, 2009). The increased levels of GDF3 in hPSC^12p^ could therefore directly support *NANOG* activation through TGFβ/SMAD2/3 signaling. Because *GDF3* expression is in part regulated by *NANOG* (Fig. S2A and (Fischer *et al.*, 2010)), a positive feedback loop could be formed between them that maintains the pluripotent state. To test this, we inhibited the TGFβ receptors ALK4, 5 and 7 with SB-431542 at day 3 of differentiation in VUB14^12p^, beyond the point where activin A induces TGFβ signaling to generate definitive endoderm. Treatment with SB-431542 led to the elimination of residual pluripotent cells at day 8 but not to a full rescue of differentiation efficiency (Fig. 5E). This indicates that the cells failed to initiate differentiation but that the pluripotent state subsequently collapsed, leading to misspecified differentiated cells instead of residual pluripotent cells. Furthermore, knockdown of *GDF3* as from day 3 of differentiation also led to a loss of residual pluripotent cells (Fig. 5F), confirming that high GDF3 is necessary for the maintenance of pluripotency seen in residual pluripotent cells and that it is likely acting through activation of TGFβ signaling.

Together, we propose a model for the formation of residual pluripotent cells during hepatoblast induction of hPSC^12p^. Excess NANOG attenuates WNT signaling through direct physical interaction with TCF7, inhibiting β-catenin induced differentiation, and by maintaining the pluripotency network. This is further supported by GDF3’s BMP inhibitory capacity. The delayed onset of WNT signaling and BMP inhibiton leads to the bifurcation of hPSC^12p^ into two separate subpopulations in a deterministic manner depending on their position within the cell cycle. S-phase cells more quickly activate WNT signaling and initiate differentiation while slower activation in G1 cells instead leads to the formation of residual pluripotent cells. Pluripotency is then maintained by a positive feedback loop through GDF3-induced TGFβ signaling, which directly maintains *NANOG* expression, which in turn maintains *GDF3* expression (Fig. 6).

**Figure 6.**
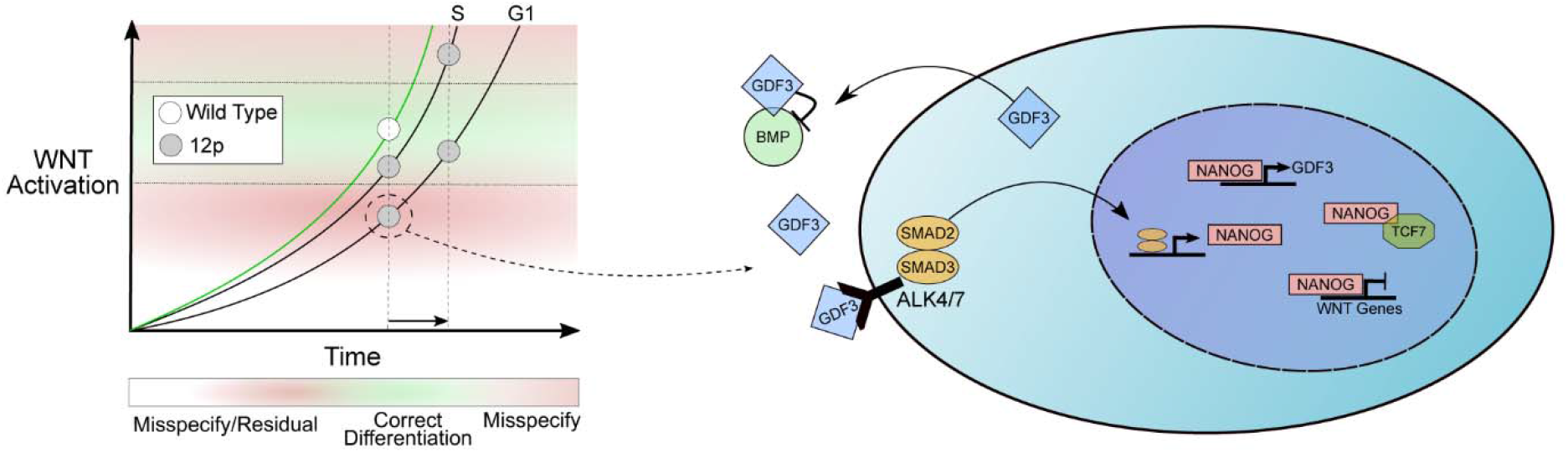
Schematic overview of proposed mechanism of residual pluripotent cell formation: hPSC^12p^ have a delayed activation of WNT as a result of NANOG overexpression, which results in cells that are in G1 to fail to reach the activation threshold needed to initiate differentiation, instead maintaining pluripotency. Continued overexpression of *NANOG* and *GDF3* create a positive feedback loop that maintains the residual pluripotent cells in an undifferentiated state through GDF3 induced activin/nodal signaling. The residual pluripotent state is further maintained by inhibition of BMP and WNT by GDF3 and NANOG, respectively.

## Discussion

To date, most work on chromosomally aberrant hPSC has focused on their impact in the undifferentiated state, dissecting the mechanisms of culture takeover and the extent and causes of their genetic instability. The impact of recurrent CNVs on differentiation is less well understood but may have an important impact on the suitability of hPSC as a research model or in clinical applications. We present here the first systematic evaluation of the impact on differentiation of segmental gains of chromosome 12p13.31, one of the most recurrent CNVs in hPSC. Using three isogenic hPSC cell lines, we identified a general reduction in differentiation capacity towards progenitors of all three germ layers, and more importantly, a mechanism that maintains a sub-population of hPSC with a gain of 12p13.31 in a stable residual pluripotent state during hepatoblast induction. We show that overexpression of *NANOG* results in a failure to induce mesendoderm differentiation by attenuating the response of hPSC^12p^ to the factors inducing the activation of WNT signaling. This effect is compounded by GDF3’s BMP inhibition. The initial failure to dissolve the pluripotent state is followed by a positive feedback loop between these two genes that continuously maintains the cells in an undifferentiated state during hepatoblast specification culture conditions (Fig. 6).

Our results further elucidate the mechanism by which *NANOG* interferes with the onset of differentiation. *NANOG* maintains the pluripotent state by blocking FGF-driven differentiation and also limiting TGFβ-induced SMAD2/3 signaling, maintaining pluripotency instead of inducing differentiation (Vallier *et al.*, 2009). However, by introducing pro-differentiation factors such as WNT, the context of TGFβ signaling changes, shifting the cell’s fate towards differentiation instead. The full mechanism of how this occurs is not yet fully understood. Here we present multiple lines of evidence that directly implicate NANOG in delaying differentiation by interfering with the activation of the WNT pathway. *NANOG* knockdown rescues decreased WNT activation seen in hPSC^12p^ during the first 24h of differentiation. This delay is due, at least in part, to NANOG’s direct physical interaction with TCF7, likely preventing it’s binding to β-catenin as previously described (He *et al.*, 2020). In line with this, single-cell RNA sequencing reveals reduced activation of WNT and mesendoderm genes in the ancestors of residual pluripotent cells during the first 24h of hepatoblast specification. It has previously been shown that a lack of WNT signaling delays mesendoderm induction (Funa *et al.*, 2015) and a recent report from our lab has demonstrated that high endogenous *NANOG* expression interferes with WNT-induced differentiation towards definitive endoderm (Dziedzicka *et al.*, 2021). Together these reports are in agreement with our findings that a NANOG-induced interference of WNT signaling delays the onset of differentiation. In hPSC^12p^ this delay in differentiation is further amplified by inhibition of endogenous BMP signaling, which is crucial for definitive endoderm specification, as a result of excess GDF3. Interference in both BMP and WNT signaling likely accounts for the reduced differentiation capacity seen across the cell types tested here.

During hepatoblast specification of hPSC^12p^, cells that do not initiate differentiation instead form a population of OCT4^+^ residual pluripotent cells. Our results show that overexpression of *NANOG* and *GDF3* together sustains pluripotency in these cells. Knockdown of either gene abrogates the formation of the residual pluripotent cells, however, knockdown of *GDF3* still leads to a heterogeneous differentiation, strengthening our findings on *NANOG’s* role in interfering with early mesendoderm specification. Importantly, we demonstrate that without treatment with CHIR99021, hPSC^12p^ fail to form residual pluripotent cells, indicating that activation of WNT signaling is crucial for entry into this state. WNT signaling has been implicated in the maintenance of pluripotency in hPSC (Sato *et al.*, 2004; Tsutsui *et al.*, 2011), however, it is unclear how WNT feeds into the pluripotency network in hPSC, and long-term WNT signaling is not suitable for maintenance of pluripotency (Dravid *et al.*, 2005). Several reports have demonstrated that short term or low induction of WNT leads to an upregulation or maintenance of *NANOG* expression (Singh *et al.*, 2012; Huang *et al.*, 2015). Consistent with this, it has been demonstrated that WNT/CHIR induced β-catenin stabilization can maintain the pluripotent state only when β-catenin is prevented from binding to the TCFs and inducing differentiation in the nucleus (Kim *et al.*, 2013). Our findings indicate that NANOG physically interacts with TCF7 and would thereby prevent β-catenin from inducing differentiation. Remaining cytoplasmic could then lead to a temporary boost in the pluripotency network as previously described (Huang *et al.*, 2015). Under normal circumstances, the NANOG-TCF7 interaction would be overcome quickly and WNT-induced differentiation would initiate, however, in hPSC^12p^, overexpressed NANOG maintains this interaction for longer, delaying the onset of differentiation, and driving cells into the residual pluripotent state.

Our results demonstrate that formation of residual pluripotent cells is cell cycle dependent, but contrary to previously published work (Sela *et al.*, 2012; Pauklin and Vallier, 2013), we observed an improvement in differentiation when cells were synchronized to S, and not G1. We propose that due to the overexpression of *NANOG,* G1 synchronized hPSC^12p^ do not have sufficient time to activate WNT signaling beyond the threshold needed to initiate differentiation. Cells that are in S/G2/M have previously been shown to be refractory to differentiation, and preferentially maintain pluripotency (Gonzales *et al.*, 2015). In our model, G1-enriched hPSC^12p^ enter into S-phase before the initiation of differentiation. Upon cycling back into G1, where they are again poised to begin differentiation, WNT signaling becomes active but fails to reach the threshold needed to initiate differentiation and instead will promote maintenance of pluripotency. This is supported by the delay in the initial onset of WNT signaling seen in G1 synchronized compared to S synchronized hPSC^12p^, and by the fact that extended CHIR exposure, and thus elongation of the activation window, pushes the cells out of pluripotency. In contrast, S synchronized hPSC^12p^ begin to activate WNT after approximately 8h, which in addition to the 8h release from nocodazole, coincides with the reported cycling length in hPSC of 16-20h (Becker *et al.*, 2006) and indicates the cells begin upregulating WNT only upon entering back into G1. S-synchronized cells would then have sufficient time to activate WNT signaling to a level needed to initiate differentiation, overcoming the attenuating effects of *NANOG* overexpression. This model would reconcile the need for hPSC to be in G1 to properly differentiate, in agreement with previous findings.

In summary, we describe a mechanism that implicates *NANOG* and *GDF3* in the formation of residual pluripotent cells during hepatoblast specification and a general reduction in differentiation capacity resulting from gains of 12p13.31 in hPSC. Recent work from our lab has also described a reduction in neuroectoderm capacity in hPSC with gains of 20q11.21 (Markouli *et al.*, 2019), suggesting that the influence of CNVs on hPSC differentiation is more prevalent than current literature indicates. The smallest known gain of 20q.11.21 is less than 1Mb in size, and we show here gains of 12p13.31 as small as 2Mb, both falling below the detection threshold of typically used banding techniques. Furthermore, both gains have also been identified as low level mosaics in hPSC (Jacobs *et al.*, 2014; Keller *et al.*, 2019). Given the impact of gains of 12p13.31 on hPSC differentiation, this raises the possibility of unidentified mosaic populations interfering with differentiation outcomes by creating heterogenous cell populations, potentially impacting research outcomes. The formation and long-term maintenance of residual pluripotent cells in differentiation promoting conditions also raises serious concerns about the tumorigenic risk of hPSC-derived cell products. Gains of chromosome 12 are strongly associated with numerous cancer and malignant cell types (Atkin and Baker, 1982; Rodriguez *et al.*, 2003; Draper *et al.*, 2004; Andrews *et al.*, 2005; Korkola *et al.*, 2006) and teratomas formed by trisomy 12 hPSC have been shown to have neoplastic properties, more closely resembling teratocarcinomas, from which undifferentiated cells can be isolated (Werbowetski-Ogilvie *et al.*, 2009; Ben-David *et al.*, 2014). It is unclear if segmental gains pose the same risk of neoplastic progression, however, our findings strongly suggest a mechanistic link to the undifferentiated cells recovered from them. As residual pluripotent cells remain one of the most important safety hurdles to overcome in hPSC-based cell therapy (Yamanaka, 2020), further work is needed to understand what risk gains of 12p13.31 pose in a clinical setting.

### Limitations of the study

While we demonstrate a need for WNT activation for the formation of residual pluripotent cells, it is unclear how WNT drives this process. As sustained CHIR99021 exposure pushes the cells out of the pluripotent state, a yet unknown mechanism temporarily drives entry into the residual pluripotent state, likely in a temporal manner. Similarly, we were unable to identify a significant difference between the residual and normal pluripotent states beyond the activation of OXPHOS. Our single cell RNAseq data suggests the cells are largely similar, though the activation of cancer-associated genes indicates that important unidentified differences may exist. Lastly, we identified a generally reduced differentiation capacity in hPSC^12p^ towards neuroectoderm and myoblasts, without the presence of residual pluripotent cells. However, we did not investigate the specific mechanisms underpinning this reduced differentiation capacity, nor is it clear what impact this will have on more mature cell types.

## Acknowledgements

The authors would like to thank their colleagues from the human embryonic stem cell lab for the derivation of lines used in this study, the BENE lab of the VUB for their assistance with the 10X library preparation and the BRIGHTcore of the UZ Brussels for assistance with all sequencing performed. We would also like to thank Prof. Dr. Frederic Lluis Viñas of the Stem Cell Institute Leuven at the KU Leuven for his advice and assistance with the WNT reporter line and signaling pathway. This work was supported by the Methusalem Grant of the Research Council of the VUB under KS. AK, NK, ECD and DD are/were all doctoral fellows at the Fonds Wetenschappelijk Onderzoek. CM was a doctoral fellow supported by the Instituut voor Innovatie doorWetenschap en Technologie.

## Author contributions

AK performed all experiments unless otherwise stated, experiment analysis and wrote the manuscript. YL and ECD performed bioinformatic processing and assisted with bioinformatic analysis. NK generated RGB and WNT reporter lines. DD and CM assisted with differentiations. KS proofread the manuscript. MG designed and supervised experimental work and proofread the paper. CS designed and supervised experimental work and co-wrote the paper.

## Experimental Procedures

### hESC culture, characterization reporter line generation

VUB14, VUB17 and VUB19 (VUB19_DM1) were derived from human embryos donated for research and characterized as previously described (Mateizel *et al.*, 2006, 2008) and are registered in the EU hPSC registry (https://hpscreg.eu/). Cells were routinely cultured in 6-well dishes at 37°C in 5% CO_2_ and atmospheric O_2_ on laminin-521 (Biolamina) at a concentration of 10μg/mL in Nutristem hPSC XF Medium (Biological Industries) containing 100U/mL penicillin/Streptomycin (Thermo Fisher Scientific). Medium was changed daily. Cells were passaged as single cells at 70-90% confluency using TrypLE Express (Thermo Fisher Scientific) and split at a ratio of 1:5 to 1:20, as needed. The genetic content of the cells was evaluated by shallow whole genome sequencing as previously described (Bayindir *et al.*, 2015). Bulks of chromosomally balanced and 12p13.31 hPSC were then generated and stored for use in all experiments.

### Reporter line generation

The RGB vectors were kindly provided by Kristoffer Riecken (http://www.lentigo-vectors.de/index.htm) (Weber *et al.*, 2011) and the WNT reporter 7TGP was a gift from Roel Nusse (Addgene plasmid # 24305; http://n2t.net/addgene:24305; RRID:Addgene_24305) (Fuerer and Nusse, 2010). Briefly, lentiviruses were produced in HEK 293T cells by transfecting plasmids for VSV.G, gag-pol and the plasmid of interest in PEI (2μg per μg of DNA; Polysciences Inc) and Opti-MEM (Thermo Fisher Scientific) for 4h. The transfection cocktail was then replaced with complete medium and the lentivirus-containing supernatant was harvested 48h-72h later and stored in aliquots at −80°C. One day before transduction, hPSC were seeded at a density of 50,000 cells per well of a 6well plate. Cells were then transduced in a transduction cocktail with 1:1000 protamine sulfate (LEO Pharma; 10mg/mL) and a 50:50 mix of Nutristem and lentivirus-containing medium. Transduction cocktail was added to each well of a 6-well plate and incubated for 4h. Cells were then washed with PBS 5x before adding fresh Nutristem medium. 24h later cells were again washed with PBS 5x before being selected for by either puromycin, FACS or both.

### Directed differentiation

Hepatoblast differentiation was adapted from Cameron et al., previously optimized for laminin-521 (Cameron *et al.*, 2015). Briefly, cells were seeded at 50,000 cells/cm^2^ 1-2 days before the onset of differentiation. Differentiation was initiated when cells reached 30-40% confluency. Medium was changed from Nutristem to RPMI 1640 with Glutamax supplemented with 0.5% B27 (Thermo Fisher Scientific), 100ng/mL activin A (R&D systems) and 3μM CHIR99021 (Stem Cell Technologies) for 24h, followed by an additional 24h in the same medium in the absence of CHIR. Medium was then changed to hepatoblast medium consisting of KnockOut DMEM with 20% KnockOut Serum Replacement, 1% MEM NEAA, 0.5% Glutamax, 100U/mL penicillin/streptomycin (Thermo Fisher Scientific), 1% DMSO and 0.1 mM β-mercaptoethanol (Sigma-Aldrich). Medium was changed daily until day 8. Neurectoderm differentiation was performed as previously described (Markouli *et al.*, 2019). Briefly, cells were seeded on laminin-521 coated dishes and grown until 90-95% confluent. Medium was changed from Nutristem to neurectoderm specification medium consisting of KnockOut DMEM with 10% KnockOut Serum Replacement (Thermo Fisher Scientific), supplemented with 500ng/mL human recombinant noggin (R&D Systems) and 10μM SB431542 (Torcis). Medium was changed daily for 8 days. Myoblast differentiation was adapted from (van der Wal *et al.*, 2017). 50,000 cells were seeded per well in laminin-521 coated 24-well plates, the following day medium was changed daily for 48h to DMEM/F12 containing 1X ITS-X, 1X Penicillin-Streptomycin-Glutamine (Thermo Fisher Scientific) and 10μM CHIR99021 (Stem Cell Technologies). After 48h, CHIR99021 was substituted for 20ng/mL FGF2 (Peprotech) and changed daily until day 12.

### Immunostaining and live imaging

For immunostaining, cells were fixed in 3.7% formaldehyde, permeabilized in 0.1% Triton X (Sigma-Aldrich) and blocked with 10% fetal bovine serum (Thermo Fisher Scientific). Primary antibodies were incubated overnight at 4°C and secondary antibodies were incubated for 1h at room temperature. Imaging was performed on a LSM800 confocal microscope (Carl Zeiss). Live imaging was performed using the LSM800 with incubator stage. Images were captured in 10-20m intervals for 24h. A list of antibodies used in this study can be found in supplemental table 4.

### qRT-PCR

Total RNA was extracted using RNeasy Mini and Micro kits (Qiagen) and converted to cDNA with First-Strand cDNA Synthesis Kit (Cytiva). Real-time qPCR was performed using qPCR Mastermix Plus – Low ROX (Eurogentec) on a ViiA 7 thermocycler (Thermo Fisher Scientific) using the standard cycling protocol. Samples were run in triplicate using GUSB and UBC as endogenous controls. A list of probes and assays used in this study can be found in supplemental table 5.

### Single cell sequencing

Cells were differentiated simultaneously and harvested at 12h, 24h, 48h, 72h and day 8 of hepatic differentiation. Briefly, cells were washed 3X in DPBS, dissociated using TrypLE (Thermo Fisher Scientific), strained through a 20μm cell strainer (pluriSelect) and centrifuged for 5min at 200g. Cells were then resuspended in culture medium and counted using a Tali Image Cytometer (Thermo Fisher Scientific), diluted to a concentration of 1000 cells/μL and taken for library preparation. Single cell libraries were generated using the 10X Chromium Controller (10X Genomics) according to manufacturer’s instructions. Approximately 1000 cells were sequenced per time point to minimize the occurrence of duplets. Sequencing was performed on a NovaSeq (Illumina) with 50 million reads per sample.

### Single-cell RNA-seq processing and Bioinformatics

The BCL files were demultiplexed using the CellRanger mkfastq pipeline. The reads alignment, filtering barcode counting and UMI counting were performed using CellRanger 3.1.0 (10X Genomics). The reads were aligned to the Genome Reference Consortium Build 38. The different 10X genomics runs were aggregated using the CellRanger aggr pipeline. The processed scRNA-seq data were analyzed using R (3.6.3) (https://www.R-project.org/) with the Seurat library v3 (Stuart et al., 2019). The cells included in the analysis had nFeature counts ∈] 500;8000 [and mitochondrial counts between ∈] 1%;20%[. The data were scaled. The data were normalized using the LogNormalize function of Seurat. We regressed the cell-cell variation on the nCount_RNA, percentage of mitochondrial gene content and percentage of ribosomal gene content. Differential gene expression analysis between the cells among the clusters was determined using the FindMarkers function with the default parameter using Wilcoxon rank sum test. Genes were ranked by sign(log2FC)*(-log(p-value)) gene set enrichment analysis (GSEA) was performed using GSEA function embedded in the clusterProfiler package and H hallmark gene sets from MSigDB database. Ingenuity Pathway Analysis canonical pathway and upstream transcription factor prediction was performed using the 668 significantly differentially expressed genes, only predictions based on >10 genes were considered. The cell cycle analysis was conducted using Cell Cycle Scoring function as described in (Tirosh *et al.*, 2016). Using default settings, cells were assigned a cell-cycle phase based on the expression levels of S and G2/M phase markers, cells highly expressing S or G2/M-phase markers were assigned to the relevant phase and cells expressing neither were assigned to G1.

### siRNA and Plasmid transfection

SMARTpool siRNA’s were purchased from Horizon Discovery and transfected at a concentration of 50nM in Nutristem medium in the absence of penicillin/Streptomycin with RNAiMAX (Thermo Fisher Scientific), according to the manufacturer’s instructions. hPSC were transfected for either 24h prior to the onset of hepatoblast differentiation, or on days 3, 5 and 7 during differentiation. Seeding density was adjusted to account for cell growth prior to the onset of differentiation. pENTR221-TCF7 was a gift from Peter Jon Nelson (Addgene plasmid # 79498; http://n2t.net/addgene:79498; RRID:Addgene_79498)(Jäckel *et al.*, 2016) and FLAG-NANOG was purchased from Sino Biological Inc. Transfection was performed with Lipofectamine 3000 (Thermo Fisher Scientific) according to the manufacturer’s instructions, with an input of 5μg per well of a 6 well plate. Cells were transfected for 24h and allowed to grow for an additional 24h before harvesting.

### Cell synchronization

hPSC were treated with 5ng/mL nocodazole (Sigma-Aldrich) for 16h to synchronize to G2/M as previously described (Yiangou *et al.*, 2019). Cells were then allowed to recover for 2h or 8h prior to the initiation of differentiation to enrich in G1 or S. Cell cycle position was confirmed by a 30m EdU (Thermo Fisher Scientific) pulse for S-phase cells and HH3 stain for G2/M.

### Co-IP and Western Blot

For protein isolation, cells were washed 3 times with PBS and harvested with Trypsin-EDTA (Thermo Fisher) for 5m at 37°. Trypsin-EDTA was inactivated with an equal volume of medium and pelleted by centrifugation at 300g for 5min. Protein isolation and IP was performed using the Pierce Crosslink Magnetic IP/Co-IP Kit according to manufacturer’s instruction. Protein was quantified using a Qubit protein assay kit (Thermo Fisher Scientific). FLAG-NANOG was captured using anti-FLAG antibody (Sigma-Aldrich). Samples were prepared for Western Blot by diluting 8μg protein 1:4 with loading buffer (1:10 β-mercaptoethanol (Sigma-Aldrich) and 9:10 laemmli (Bio-Rad)). Sample-buffer mixtures were heated to 95°C for 5min and kept on ice before loading into precast TGX gels and run in 1X T/G/S buffer in a Criterion Vertical Electrophoresis Cell. Precision Plus Protein ladder was used to identify protein size. Protein was transferred to a nitrocellulose membrane using the Trans-Blot Turbo system (all from Bio-Rad). The nitrocellulose membrane was blocked in 5% non-fat milk for 1h, washed with 0.1% PBS-Tween (Sigma-Aldrich) and incubated with the primary antibody on a rotating platform in 5% FBS/0.1% PBS-Tween overnight at 4°C. The following day samples were washed 3X 10 minutes with 0.1% PBS-Tween and the secondary antibodies were incubated on a rotating platform for 1h at room temperature 5% FBS/0.1% PBS-Tween and washed again 3X 10 minutes with 0.1% PBS-Tween. Samples were imaged on an Odyssey FC Imaging System (LI-COR). A list of antibodies used in this study can be found in supplemental information.

