## Supplementary Figures for "Gains of 12p13.31 delay WNT-mediated initiation of hPSC differentiation and promote residual pluripotency in a cell cycle dependent manner"

### Human pluripotent stem cells with gains of 12p13.31 result in a failure to exit pluripotency upon directed hepatic differentiation

*Alexander Keller, Yingnan Lei, Nuša Krivec, Edouard Couvreu de Deckersberg, Dominika Dziedzicka, Christina Markouli, Karen Sermon, Mieke Geens*, Claudia Spits*[1]*

**joint last authorship*

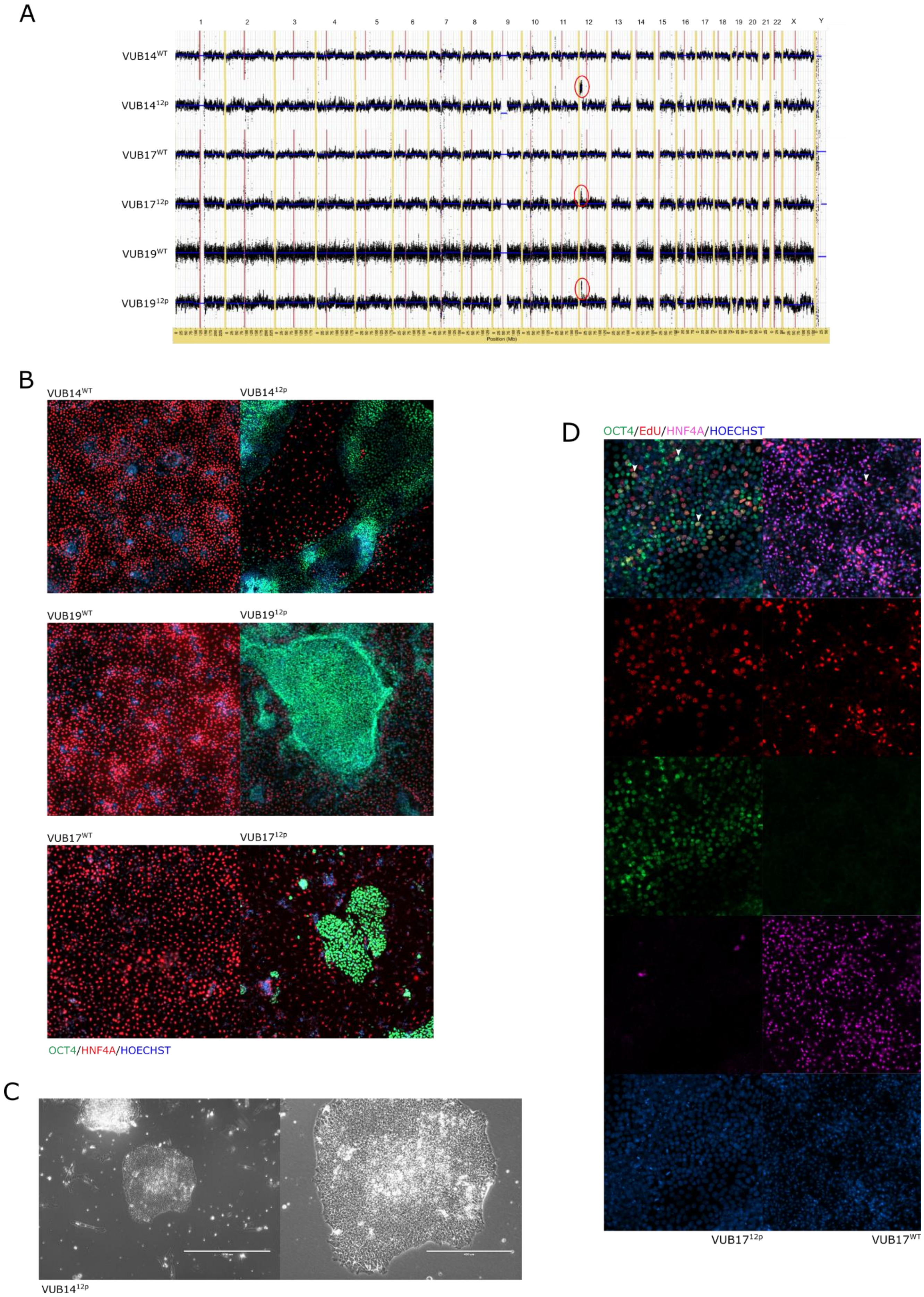
A
1 2 3 4 5 6 7 8 9 10 11 12 13 14 15 16 17 18 19 20 21 22 X Y
VUB14WT
VUB1412p
VUB17WT
VUB1712p
VUB19WT
VUB1912p
Position (Mb)
B
VUB14WT
VUB1412p
VUB19WT
VUB1912p
VUB17WT
VUB1712p
OCT4/HNF4A/HOECHST
C
VUB1412p
D
OCT4/EdU/HNF4A/HOECHST
VUB1712p
VUB17WT

**Figure S1. Gain of 12p13.31 impairs exit of pluripotency in a subpopulation of hPSC during hepatoblast differentiation. Related to Figure 1:** A) Shallow sequencing of all cell lines used in this study, gains highlighted by red circles B) Immunostaining for HNF4A (red) and OCT4 (green) after 8 day hepatoblast induction in isogenic pairs of VUB14, VUB17 and VUB19. C) Phase contrast image of $VUB14^{12p}$ residual pluripotent cell colony isolated after 22 days of hepatoblast differentiation. D) Representative immunostaining of $VUB17^{WT}$ and $VUB17^{12p}$ after EdU incorporation, only cells double positive for EdU and either HNF4A or OCT4, respectively, were counted.

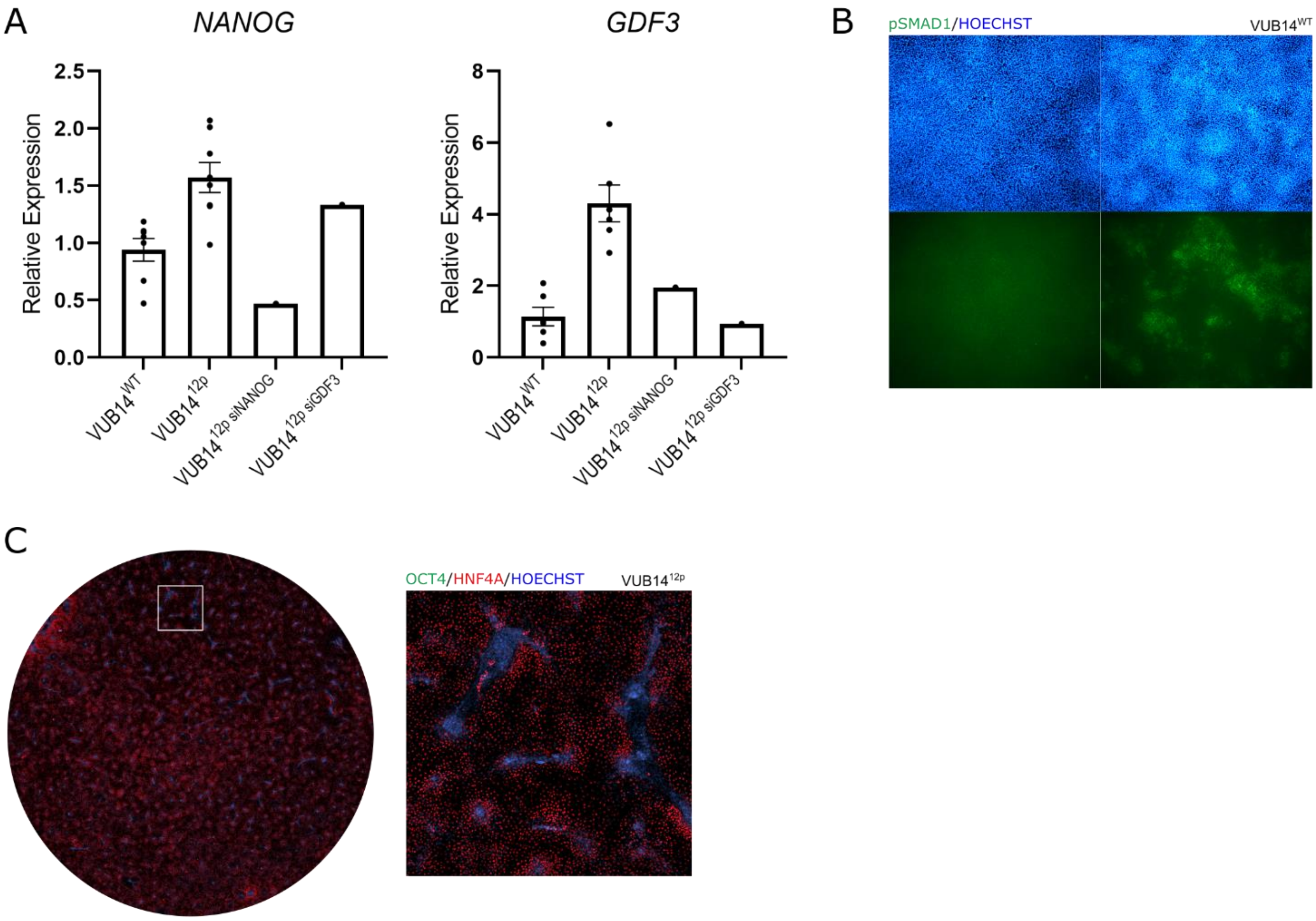


**Figure S2. Overexpression of NANOG and GDF3 induce entry into residual pluripotent state in hPSC$^{12p}$. Related to Figure 3:** A) Gene expression levels of GDF3 and NANOG in undifferentiated VUB14$^{12p}$ after siRNA knockdown (siRNA conditions n = 1) B) pSMAD1 immunostaining of VUB14$^{WT}$ after 24h of hepatoblast induction. C) OCT4 (green) and HNF4A (red) Immunostaining of day 8 hepatoblasts of VUB14$^{12p}$ with addition of 8ng/mL BMP4 in the first 24h of differentiation.

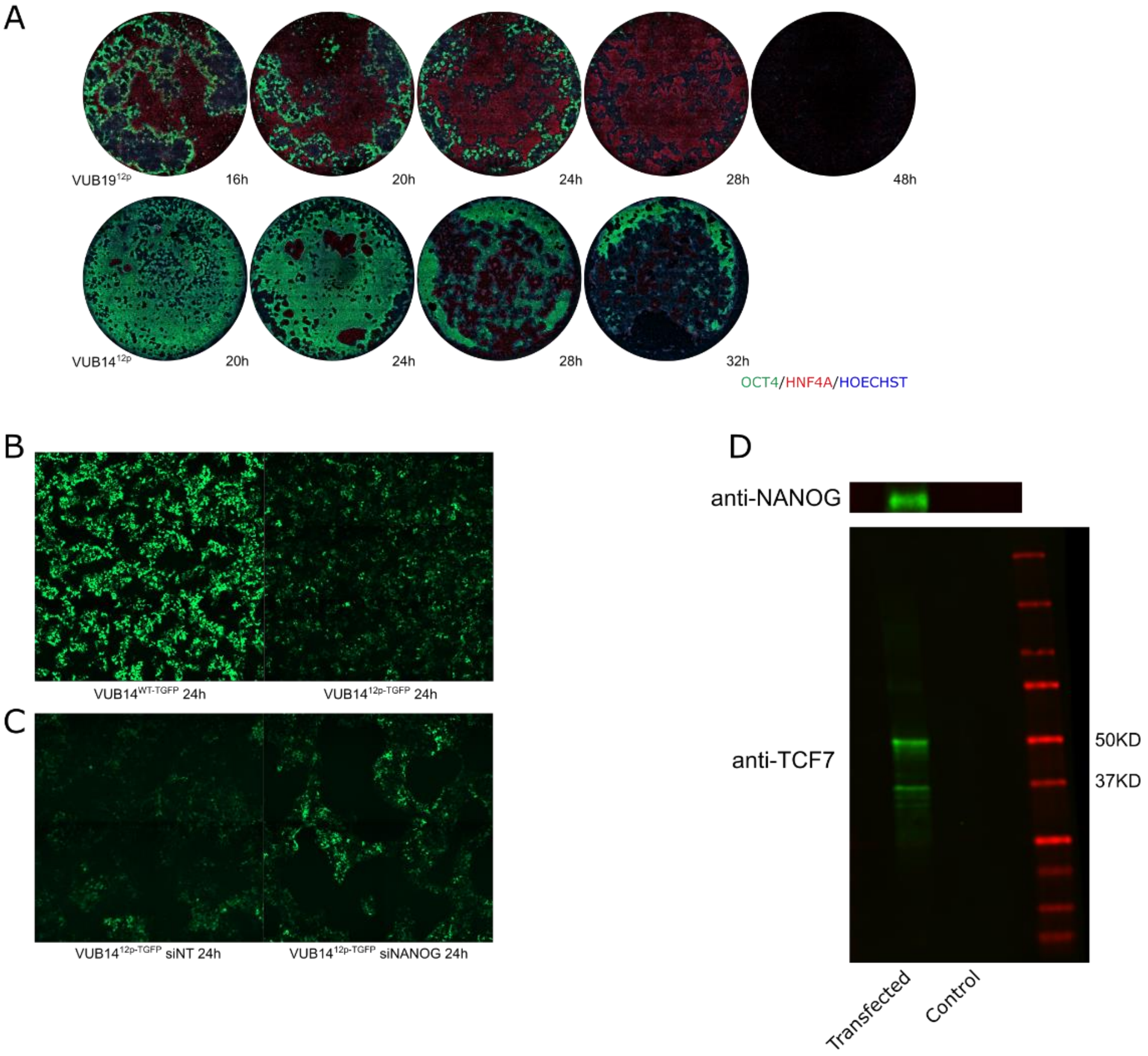


**Figure S3. hPSC$^{12p}$ have a delayed onset WNT activation as a result of NANOG overexpression. Related to Figure 4:** A) OCT4 (green) and HNF4A (red) immunofluorescent stain of day 8 hepatoblasts of VUB14$^{12p}$ and VUB19$^{12p}$ after extending CHIR99021 treatment in 4h increments. B-C) Live cell image of VUB14$^{WT\text{-}TGFP}$ and VUB14$^{12p\text{-}TGFP}$ after 24h hepatoblast induction under control and si*NANOG* conditions. D) Co-IP capture of FLAG-tagged NANOG. Western blot of captured NANOG (green) and for TCF7 (green). Lane 1 293T cells transfected with FLAG-NANOG and HA-TCF7, lane 2 non-transfected control 293T cells. Two separate isoforms of TCF7 are detected at 50 and 37 kilodaltons. Input of 8ug per lane.

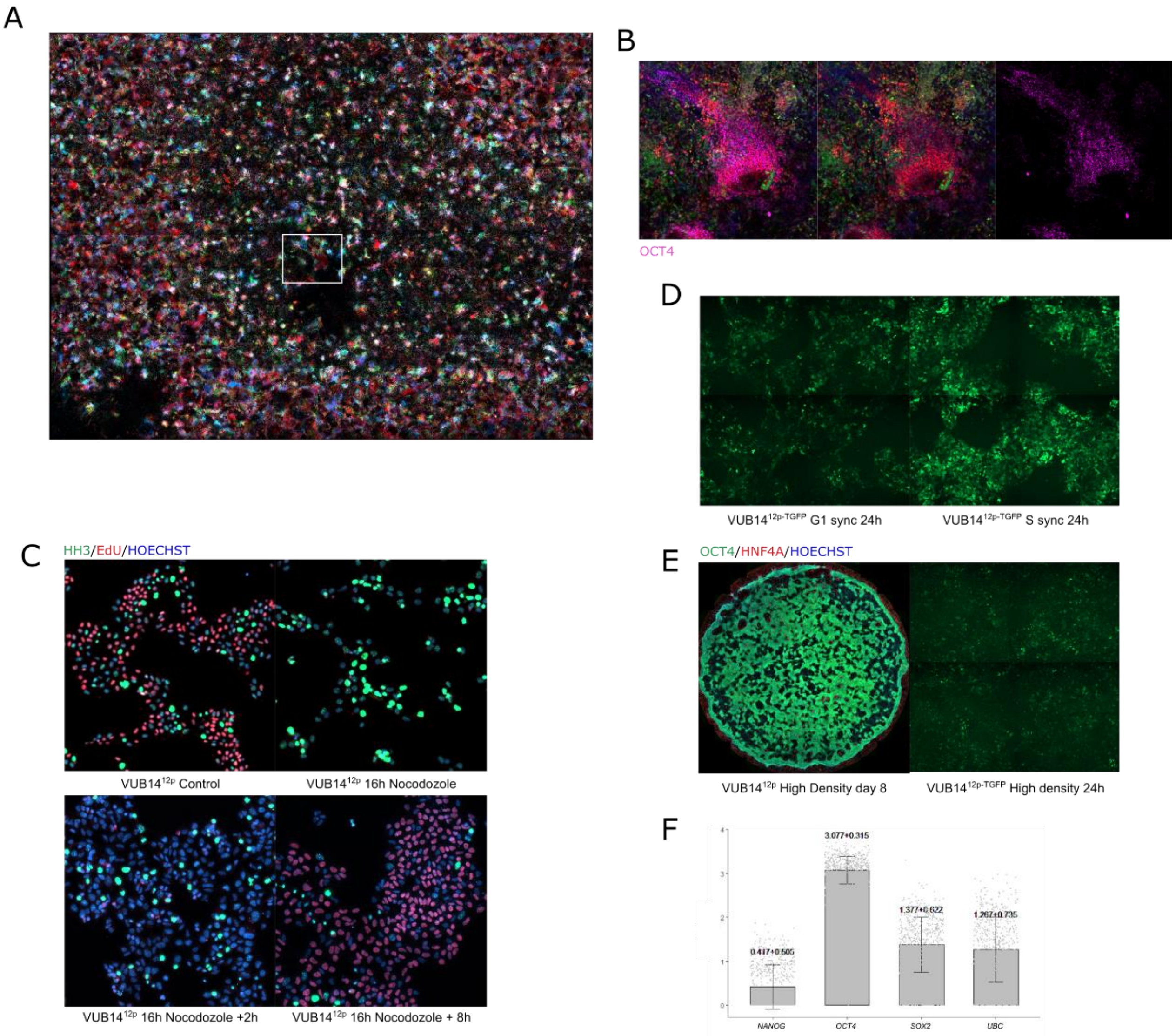


**Figure S4. Delayed WNT signaling in hPSC$^{12p}$ is cell cycle dependent. Related to Figure 5:** A) Whole dish image of VUB19$^{12p\text{-}RGB}$ demonstrating broad diversity in colors across the dish and B) an example of an OCT4$^{+}$ colony. C) HH3 (green) and EdU (red) immunofluorescent stain of VUB14$^{12p}$ under control and nocodazole treatment, and after 2h and 8h release and enrichment in G1 and S, respectively. Cells synchronized to G2/M are positive for HH3, cells in G1 are double negative for HH3 and EdU, cells in S are positive for EdU. D) Live cell image of VUB14$^{WT\text{-}TGFP}$ and VUB14$^{12p\text{-}TGFP}$ after 24h hepatoblast induction synchronized to G1 and S. E) OCT4 (green) and HNF4A (red) immunofluorescent stain of VUB14$^{12p}$ at day 8 of hepatoblast differentiation and live cell image of VUB14$^{12p\text{-}TGFP}$ after 24h hepatoblast induction after a high density (80+% confluency) initiation of differentiation. F) Distribution of single cell expression of *NANOG*, *OCT4*, *SOX2* and *UBC* in undifferentiated VUB14$^{12p}$ from cluster 1 cells (Fig. 2a).

**Supplemental Table 1. List of genes driving the GSEA prediction for positive enriched OXPHOS**

| HALLMARK_OXIDATIVE_PHOSPHORYLATION | FC | FDR |
|---|---|---|
| ATP5F1E | 1.11925029 | 1.67E-76 |
| ATP5MC1 | 1.01607294 | 4.04E-56 |
| ATP5MC2 | 0.71671717 | 1.16E-38 |
| ATP5ME | 1.94243656 | 1.17E-134 |
| ATP5MF | 1.71709248 | 3.94E-122 |
| ATP5MG | 1.23047227 | 1.48E-91 |
| ATP5PD | 1.16891708 | 6.56E-76 |
| ATP5PF | 1.05286887 | 1.82E-68 |
| ATP6V0B | -0.867367 | 4.64E-50 |
| ATP6V0E1 | -0.5889532 | 8.94E-26 |
| COX17 | 0.94562258 | 8.95E-70 |
| COX5B | 1.4234371 | 3.66E-103 |
| COX6A1 | 1.55314828 | 1.8E-109 |
| COX6B1 | 1.73502783 | 3E-123 |
| COX6C | 1.67928163 | 2.05E-118 |
| COX7A2 | 1.582688 | 3.2E-121 |
| COX7B | 1.61992737 | 2.32E-121 |
| COX7C | 1.77624706 | 4.64E-135 |
| COX8A | 1.35569677 | 1.27E-95 |
| CYCS | -0.9895908 | 6.23E-78 |
| GPI | 0.54371095 | 1.52E-18 |
| HCCS | -0.5123337 | 9.09E-16 |
| HSPA9 | -0.5051446 | 9.46E-14 |
| IDH1 | -0.5595 | 4.62E-19 |
| LDHA | 1.21556878 | 5.95E-60 |
| MGST3 | 0.53293382 | 2.54E-19 |
| MRPL15 | -0.6378257 | 1.73E-20 |
| MRPS12 | -0.5227221 | 5.64E-20 |
| MRPS15 | -0.807033 | 1.04E-44 |
| NDUFA1 | 1.77933462 | 8.52E-130 |
| NDUFA2 | 1.27885144 | 3.39E-90 |
| NDUFA3 | 0.94269646 | 3.77E-66 |
| NDUFA4 | 1.40025872 | 4.56E-116 |
| NDUFA6 | 0.75263672 | 9.4E-36 |
| NDUFA7 | 0.69347277 | 3.94E-42 |
| NDUFB1 | 1.19766516 | 5.2E-82 |
| NDUFB2 | 1.42877981 | 1.32E-99 |
| NDUFB3 | 0.83312918 | 2.8E-48 |
| NDUFB4 | 1.61020867 | 1.83E-106 |
| NDUFB6 | 0.77959961 | 1.58E-41 |

| | | |
|---|---|---|
| NDUFB7 | 1.08676035 | 4.2E-76 |
| NDUFC1 | 1.09688987 | 2.42E-75 |
| NDUFC2 | 0.74766125 | 2.09E-40 |
| NDUFS6 | 0.99847895 | 4.58E-76 |
| NDUFV2 | -0.5524583 | 3.67E-25 |
| POLR2F | 0.61573561 | 3.02E-26 |
| PRDX3 | -0.5024903 | 1.56E-16 |
| SLC25A4 | -0.5594247 | 3.41E-21 |
| SLC25A6 | -0.6991315 | 6.2E-61 |
| TIMM10 | 0.58283728 | 1.96E-26 |
| TIMM13 | -0.5684774 | 8.51E-31 |
| TIMM8B | 0.94084973 | 4.93E-60 |
| UQCR10 | 1.66259378 | 2.16E-115 |
| UQCR11 | 1.07957032 | 1.96E-67 |
| UQCRB | 1.05540941 | 1.27E-64 |
| UQCRH | 1.21217232 | 5.67E-113 |
| UQCRQ | 1.1957632 | 3.86E-83 |
| VDAC2 | -0.5278429 | 4.01E-27 |

**Supplemental Table 2. List of genes driving the GSEA prediction for negatively enriched G2M**

| HALLMARK_G2M_CHECKPOINT | FC | FDR |
|---|---|---|
| AMD1 | -0.8394419 | 2.27E-39 |
| CHAF1A | -0.7079135 | 9.75E-29 |
| DDX39A | -0.6321495 | 1.01E-18 |
| DKC1 | -0.6542889 | 6.25E-29 |
| HMGA1 | -0.803505 | 2.13E-101 |
| HSPA8 | -1.1488765 | 1.03E-64 |
| MARCKS | 1.24950317 | 5.71E-86 |
| RAD23B | -0.5157564 | 4.37E-16 |
| SFPQ | -0.6932559 | 1.6E-30 |
| SLC7A5 | -1.0012314 | 7.34E-52 |
| SQLE | -0.7064251 | 1.25E-31 |
| SRSF1 | -0.6235822 | 1.52E-22 |
| SRSF2 | -1.0653679 | 2.28E-82 |
| SYNCRIP | -0.7400854 | 6.24E-36 |
| UBE2S | -1.1493146 | 2.27E-78 |
| YTHDC1 | 0.63612258 | 5.83E-18 |

**Supplemental Table 3. List of genes driving the GSEA prediction for negatively enriched E2F**

| HALLMARK_E2F_TARGETS | FC | FDR |
|---|---|---|
| DCTPP1 | -1.0250486 | 2.74E-58 |
| DDX39A | -0.6321495 | 1.01E-18 |
| DUT | -0.6602899 | 2.52E-27 |
| HMGA1 | -0.803505 | 2.13E-101 |
| HMGB2 | 0.66030901 | 1.87E-19 |
| LYAR | -0.9787661 | 7.55E-54 |
| NAA38 | 0.89430252 | 5.99E-63 |
| NME1 | -0.8381661 | 1.45E-51 |
| NOP56 | -0.5176525 | 1.01E-19 |
| POP7 | -0.708475 | 4.14E-35 |
| PRKDC | 0.83581796 | 5.2E-39 |
| RAN | -0.7681983 | 7.94E-78 |
| RANBP1 | -1.0293943 | 1.96E-101 |
| RPA3 | 0.702882 | 2.15E-33 |
| SNRPB | -0.7387881 | 5.83E-57 |
| SRSF1 | -0.6235822 | 1.52E-22 |
| SRSF2 | -1.0653679 | 2.28E-82 |
| SYNCRIP | -0.7400854 | 6.24E-36 |
| TFRC | 0.61864993 | 6.56E-26 |
| UBE2S | -1.1493146 | 2.27E-78 |

**Supplemental Table 4. List of antibodies used in this study**

| Antibody | Concentration | Manufacturer | Reference |
|---|---|---|---|
| HNF4A | 1/200 | Santa Cruz | sc-374229 |
| OCT4 | 1/200 | Cell Signaling | 2840 |
| HH3 | 1/500 | abcam | ab5176 |
| FLAG | 10µg/IP | Sigma Aldrich | F3165 |
| NANOG | 1/1000 | Cell Signaling | 4903 |
| TCF7 | 1/1000 | Cell Signaling | 2203 |
| pSMAD1/5 | 1/500 | Cell Signaling | 9516 |

**Supplemental Table 5. List of TaqMan and in-house assays used in this study**

| | |
|---|---|
| OCT4 | Forward 5’-GGA-CAC-CTG-GCT-TCG-GAT-TT-3’<br>Reverse 5’-CAT-CAC-CTC-CAC-CAC-CTG-G-3’<br>Probe 6-FAM-GCC-TTC-TCG-CCC-CC-MGB |
| NANOG | Forward 5’-TGC-AAA-TGT-CTT-CTG-CTG-AGA-TG-3’<br>Reverse 5’-TCC-TGA-ATA-AGC-AGA-TCC-ATG-GA-3’<br>Probe 6-FAM-CAG-AGA-CTG-TCT-CTC-CTC-MGB |
| GDF3 | Hs00220998_m1 |
| HNF4A | Hs00604435_m1 |
| PAX6 | Hs00240871_m1 |
| PAX3 | Hs00240950_m1 |
| GUSB | Hs99999908_m1 |
| UBC | Forward 5’-CGC-AGC-CGG-GAT-TTG-3’<br>Reverse 5’-TCA-AGT-GAC-GAT-CAC-AGC-GA-3’<br>Probe 6-FAM-TCG-CAG-TTC-TTG-TTT-GTG-MGB |
